# Genotype patterns in growing solid tumors

**DOI:** 10.1101/390385

**Authors:** Mridu Nanda, Rick Durrett, U Harvard, U Duke

**Author notes:** Both authors are partially supported by NSF grant DMS 1614978 from the math biology program.

## Abstract

Over the past decade, the theory of tumor evolution has largely focused on the selective sweeps model. According to this theory, tumors evolve by a succession of clonal expansions that are initiated by driver mutations. In a 2015 analysis of colon cancer data, Sottoriva et al [34] proposed an alternative theory of tumor evolution, the so-called Big Bang model, in which one or more driver mutations are acquired by the founder gland, and the evolutionary dynamics within the expanding population are predominantly neutral. In this paper we will describe a simple mathematical model that reproduces qualitative features of the observed paatterns of genetic variability and makes quantitative predictions.

## 1 Introduction

In the multistage theory of carcinogenesis, it is thought that the sequence of “driver” mutations produces a series of selective sweeps. This theory has been confirmed by whole genome sequencing of cancer cells. Ding et al [10] have identified the clonal structure of eight relapsed acute myeloid leukemia (AML) patients. In one patient the founding clone 1 accounted for 12.74% of the tumor at the time of diagnosis. The additional mutations in clones 2 and 3 may have resulted in growth or survival advantages because they were 53.12% and 29.04% of the tumor respectively. Only 5.10% of the cells were in clone 4 indicating that it may have arisen last. However, the relapse evolved from clone 4 with the resultant clone 5 having 78 new somatic mutations compared to the sampling at day 170.

A mathematical theory has been developed for the clonal expansion model to make predictions about the level of intratumor heterogeniety [4, 14], the number of passenger (neutral) mutations in cancer cells [2, 39], and more practical questions such the evolution of resistance to treatment [21, 27, 38], and the effectiveness of combination therapy [3, 25, 28]. See [12] for an introduction to the mathematics underlying many of these applications.

The picture of successive selective sweeps has been confirmed by comparing primary tumors and their metasases [42], and by regional sequencing of breast cancer [29], glioblastoma [33], and renal carcinoma [17]. Thus it was surprising when Sottoriva et al [34] introduced and validated a ‘Big Bang’ model in which all driver mutations were present at the time of tumor initiation. They collected genetic data of various types from 349 individual tumor glands sampled from the opposite sides of 15 colorectal tumors and large adenomas. Data presented in Figure 3 of their paper shows that adenomas were characterized by mutations and copy number aberrations (CNA) that segregated between tumor sides. In contrast the majority of carcinomas exhibited the same private CNA in individual glands from different sides of the tumor. Following up on this observation, Ryser et al [31] found evidence of early abnormal cell movement in 8 of 15 invasive colorectal carcinomas (“born to be bad”) but not in four benign adenomas. For more about recent developments see the review article by Sun, Hu, and Curtis [36].

The mutation patterns found in [34] are similar to those found by Hallatschek et al [18] in a remarkable experimental paper. Two fluorescently labeled strains of *E. coli* were mixed and placed at the center of an agar plate containing a rich growth medium. The central region of the plate exhibits a dense speckled pattern reminiscent of the initial mixed population. From this ring toward the boundary of the colony, the population segregates into single colored sectors with boundaries that fluctuate. [20] and [26] have developed and analyzed models for the development of sectors in the system.

In previous work [13], we used the biased voter model [40, 7, 6] to model the microscopic tumor dynamics. The genealogies in that model are difficult to study, so we used ideas of Hallatschek and Nelson [19] to build an approximate model of the movement of genealogies. That attempt failed to reproduce observed patterns because it predicted extensive heterogeneity at small scales. We attribute that failure to continual turnover of cells in the interior of the tumor, so here we adopt a new approach in which tumor growth only occurs at the boundary. We will introduce four models: two for two-dimensional tumors and two for three dimensional tumors. In the terminology of interacting particle systems, two will use voter model dynamics, and two will use contact process dynamics. We describe the models and out results in the next four sections. A final section describes our conclusions.

## 2 Voter model *d* = 1 + 1

This model can be thought of as taking place in 1 + 1 dimensions, i.e., one space and one time. To formulate the model let *L* = {(*k,í*) ∈ Z^2^ : *í* ≥ 0, *k* + *l* is even}. It would be more natural to have a growing circular tumor, but that is awkward for computations. The approach we take here should be thought of as studying a tumor that is large enough so that when we view the dynamics at the level of individual cells the curvature of the boundary can be ignored.

To study the dynamics we suppose that at time 0 each cell has its own type, which we set equal to its first (spatial) coordinate. We add a new layer of cells at each time step. The cell at (*k, l*) is the type of the cell at (*k* – 1, *l* – 1) with probability 1/2 and the type of the cell at (*k* + 1, *l* – 1) with probability 1/2, with the choices for different cells being independent. The use of *L* may look odd but it has the desirable property that the set of sites occupied by one type is always an interval

In making pictures of simulations the fact that are no cells at (*k, l*) with *k + l* odd is annoying so we adopt the mathematically equivalent approach that at even times (*k, l*) will imitate (*k, l* – 1) or (*k* + 1, *l* – 1) with equal probability, while at odd times (*k, l*) will imitate (*k, l* –1) or (*k* – 1, *l* – 1) with equal probability.

Figure 1 shows a simulation.

**Figure 1:**
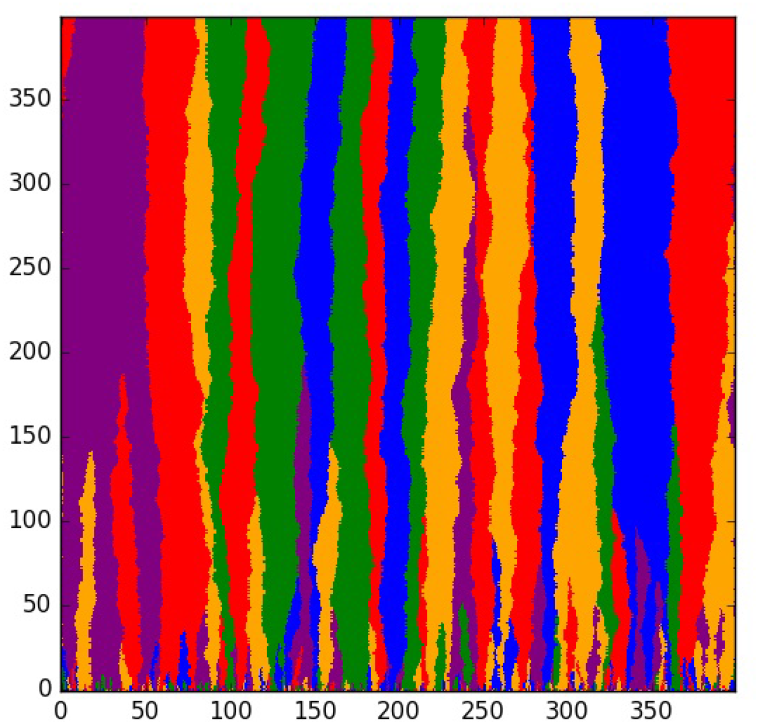
Simulation of voter model.

This system is easy to analyze because the boundaries between types are random walks. In proving the next result we will consider the model on *L*.

### Lemma 1.

*Let* 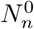 *be the number of individuals at time n whose ancestor at time 0 is at 0. Then* 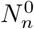 *is a lazy random walk with an absorbing state at 0 and transition probability p*(*k, k* + 1) = *p*(*k, k* – 1) = 1/4, *p*(*k, k*) = 1/2 *for k* > 0.

*Proof.* Suppose that points at −2*j*, −2*j* + 2,... − 2,0 are red at time n. If *j* ≥ 1 then the red region will surely contain − 2*j* + 1,... – 1. It will contain 1 with probability 1/2, and will contain −2*j* – 1 with probability 1/2, so if the size *k* = *j* + 1 ≥ 2 the size goes from *k* + 1 to *k, k* + 1 and *k* + 2 with probabilities 1/4, 1/2, and 1/4. Suppose now that *j* = 0, so the size *k* = 1. By looking at the four cases for the choice of parent by −1 and by 1, then the size goes to 2, 1, and 0 with probabilities 1/4, 1/2, and 1/4. □

Let *T*_0_ = min{n : 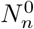 = 0}. Known results for the simple random that jumps by ±1 at each step extend easily to the lazy case with the result that

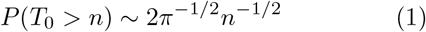

The decay of the number of regions can also be proved by using the duality between the voter model and coalescing random walk. To explain this on *L*, we draw an arrow from (*k, l*) → (*k* − 1, *l* − 1) or (*k, l*) → (*k* + 1, *l* − 1) depending on which site (*k, l*) imitated. To determine the type of *m* at time *n* we follow its sequence of ancestors (*m, n*), (*S*_1_, *n* − 1),... (*S_n_*, 0) working backwards to determine that the type is *S_n_*. The path *k* → *S_k_* is a simple random walk. Let *T_k_* be the genealogy of another site (*j, n*). If T*_l_* = *S*_l_ then the two will agree at all later times. In words, when the random walks hit they coalesce to 1. It follows from results of Bramson and Griffeath [5] that the density of coalescing random walks and hence the number types at time n decays line *cn*−^1/2^.

**Figure 2:**
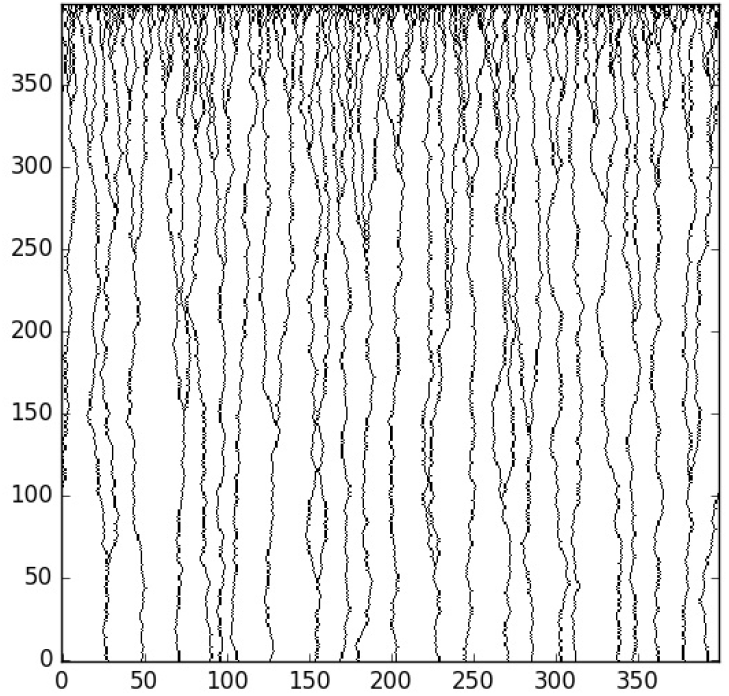
Genealogies in the voter model

**Figure 3:**
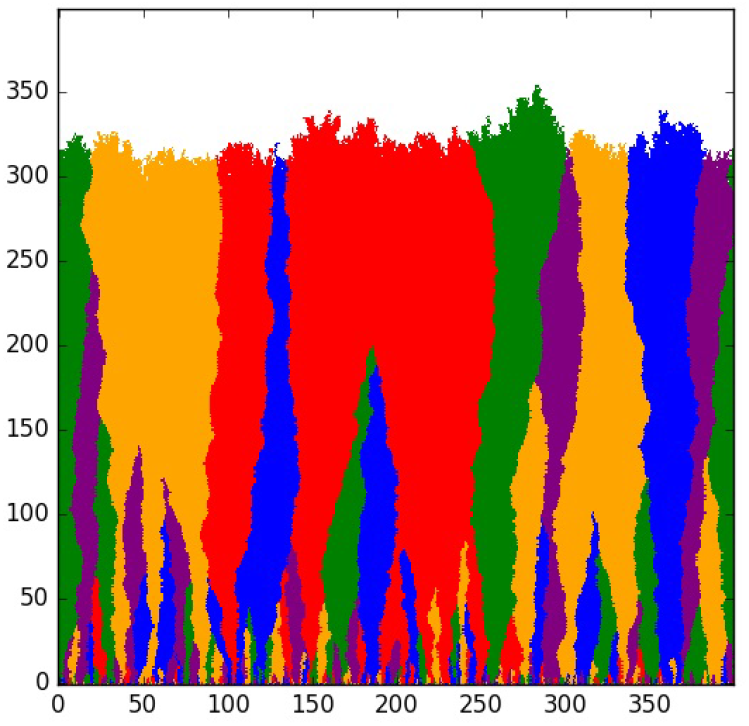
Simulation of the contact process.

Let 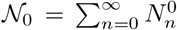 be the total number of descendents of the cell at 0 at time 0. Combining (1) with the fact that the typical distance between boundaries is 1 over the number of surviving lineages, we see that

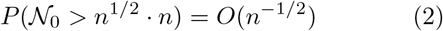

Changing variables *n* = *k*^2/3^ suggests

### Theorem 1.

*As* 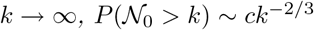.

The proof, which is hidden away in an appendix, gives a precise value for the constant and involves some interesting facts for Brownian motion.

## 3 Contact process *d*=1 + 1

Again we formulate the model on 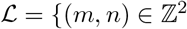 : *m* ≥ 0, *m* + *n* is even}. A particle at (*m, n*) gives birth onto (*m* + 1, *n* + 1) at rate one and onto (*m* − 1, *n* + 1) at rate one, i.e., if we let 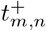 and 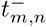 be independent exponential with mean one, then births to (*m* + 1, *n* + 1) and (*m* − 1, *n* + 1) are attempted at times 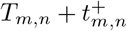 and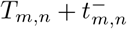, where *T_m,n_* is the time (*m, n*) is first occupied. An attempted birth succeeds if the target sites is not yet occupied. Figure 3 gives a picture of a simulation.

To be able to simulate the process efficiently, we compute the times one row at a time using

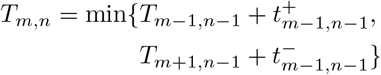

If the first time is smaller we set the type *ξ*(*m, n*) = *ξ*(*m* − 1, *n* − 1). Otherwise *ξ*(*m, n*) = *ξ*(*m* + 1, *n* − 1). These dynamics are equivalent to oriented first passage percolation on 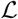. We think of 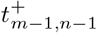 as the time it takes a fluid to traverse the edge from (*m* − 1, *n* − 1) → (*m, n*). With this interpretation *T*(*m,n*) is then the first time fluid appears at (*m, n*) if all points in 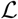 are wet at time 0. The paths that achieve the minimum time are called geodesics. They are the analogue of genealogies in the voter model.

**Figure 4:**
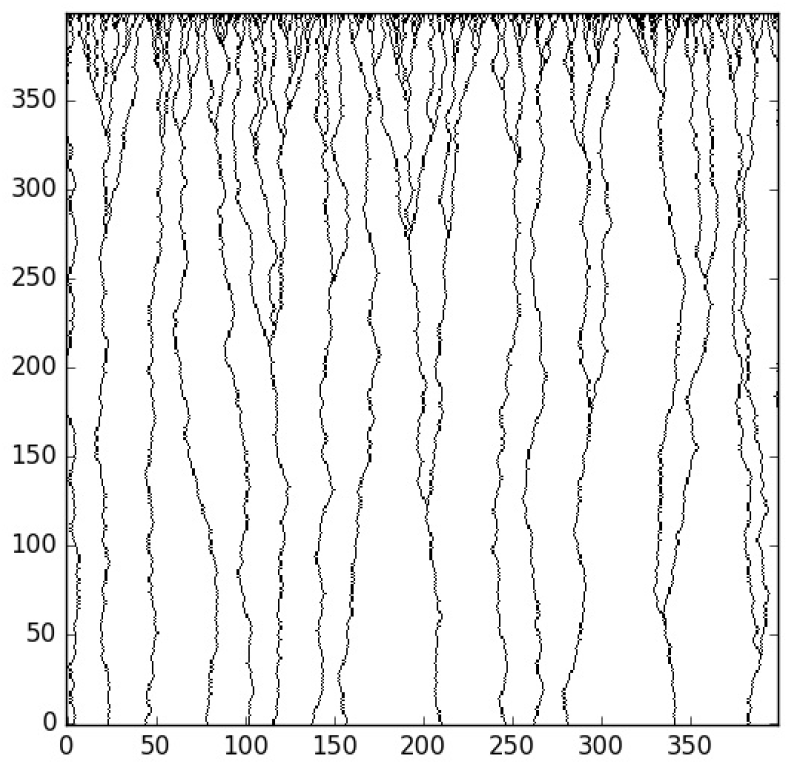
Genealogies in the contact process

There is a large literature on this and closely related process, see [1]. To explain the connection we begin by defining the growth model Eden [15] introduced in 1951. It starts with the origin occupied at time 0. At each subsequent time one vacant site adjacent to the current set of occupied sites is *ξ_n_* added. In 1973 Richardson [30] showed the set grew linearly and had an asymptotic shape. In 1986 Kardar, Parisi, and Zhang [24] studied a growing Eden cluster interface in which randomly chosen vacant sites become occupied and argued that the rescaled height converged to the solution of a stochastic partial differential now called the KPZ equation.

More closely related to our work is that of Derrida and Dickman [9] who studied an interface problem for an Eden model that starts with blue particles at (*x*, 0) with *x* < 0 and red particles at (*x*, 0) with *x* ≥ 0. They showed that on the line *y* = n the distance of the interface from the origin is of order *n*^2/3^. For more simulations see Saito and Muller-Krumbhaar [32].

Hallatschek et al [18] found similar behavior for the interfaces between different colors in their experiments with fluorescently labeled yeast strains. Statistical analysis of data showed that the wandering exponent for boundaries was *ζ* = 0.65 ± 0.05 and based on [9] concluded that *ζ* = 2/3. Figure 5 gives our results for the decrease of the number of types versus time.

**Figure 5:**
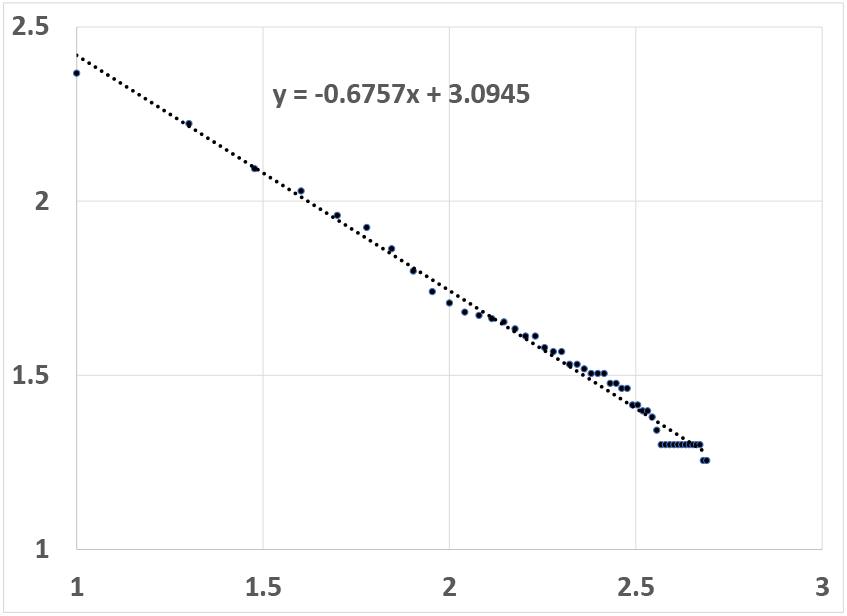
Log-log plot of the number of regions versus time in the 1 + 1 dimensional contact process. Slope is 0.6757 « 2/3.

We cannot use the theory of coalescing random walks, as we did for the voter model, but using the fact that the typical distance between boundaries is 1 over the number of surviving lineages, we see that

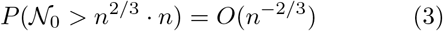

Changing variables *n* = *k*^3/5^ suggests that when *k* is large 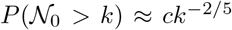. This agrees with the result found by Fusco et al [16]. See their Figure 3.

## 4 Voter model in *d* = 2 + 1

We formulate our model on 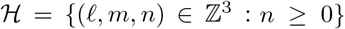. The cell at (*¿, m, n*) imitates one of five individuals from the previous time probability 1/5 each:

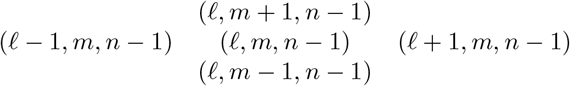

The site (*¿, m, n*) is included because otherwise there would be no interaction between the sites with *l + m + n* even and those with *l + m + n* odd. As in the voter model in 1 + 1 dimensions, we assume that at time 0 all the sites have different types which are given by their spatial location. As Figure 6 shows, the clusters of sites with the same type are small and are intermixed.

**Figure 6:**
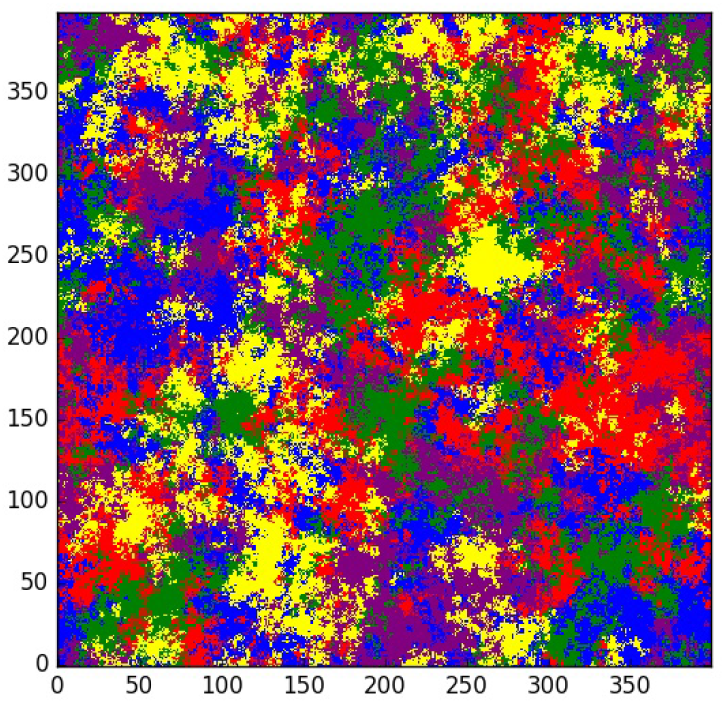
Two dimensional voter model at time 2000

To study the distribution of types in this process we again use the duality between the voter model and coalescing random walk. As in 1 + 1 dimensions we draw an arrow from (*l, m, n*) to the site in the previous generation that it imitated. To determine the state of (*x*_0_, *y*_0_, *t*) we follow the path of arrows backward to time 0. If the path ends at (*x_t_*, *y_t_*, 0) then (*x*_0_, *y*_0_, *t*) has type (*x_t_*, *y_t_*). The path (*x_s_*, *y_s_*, *t* − *s*) can be thought of as the genealogy of the individual (*x*_0_, *y_0_, *t*). The first two coordinates of the path is a random walk that jumps to the four nearest neighbors of (*x*_m_*, *y_m_*) with probability 1/5 each and stays where it is with probability 1/5.

If we consider random walks starting from two starting points (*x*_0_, *y*_0_, *t*) and (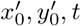) then the two genealogies agree after they hit. It is for this reason the collection of dual paths is said to be a coalescing random walk. In two dimensions random walk is recurrent, so any two genealogies will eventually hit. This implies that if *x* ≠ *y* then (*ξ_t_* (*x*) ≠ *ξ_t_*(*y*)) → 0, so if we look at the voter model on a square with side *L* then the probability we see two different types at time *t* converges to 0 as t → ∞. As the simulation in Figure 6 suggests this takes a very long time

To quantify this we begin with result of Bramson and Griffeath [5] which shows that the density of particles in a continuous time two-dimensional coalescing random walk that jumps at rate one to one of its four nearest neighbors converges to 0 like

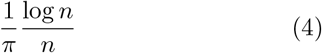

In our discrete time coalescing random walk the density is ~ *C*(log *n*)/*n*. It is not easy to compute the constant *C* for our discrete time random walk, but if we use *C* = 1/*π* we would conclude that there are an average of 13.5 types present in a box of side 400 at time 2000.

While the picture of the voter model may be pretty, the distribution of types is not what we expect in a slice through a three dimensional solid tumor. Results for random walks imply that the probability that random walks starting from adjacent sites have not coalesced by time t is ~ *c*/ log *t*. From this it follows that the number of genotypes in a *t*^1/2^ × *t*^1/2^ is of order *t*/(log *t*), so there is considerable heterogeneity at small scales. For this and more on the intricate spatial structure of the two dimensional voter model, see Cox and Griffeath [8].

## 5 Contact process in *d* = 2 + 1

Again we formulate our model on 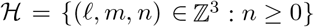. Let *z*_1_ = (− 1, 0), *z*_2_ = (0,1), *z*_3_ = (0, 0), *z*_4_ = (0, −1), and *z*_5_ = (1,0). If we let *w* = (*l, m*) and 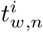, 1 ≤ *i* ≤ 5 be independent exponential with mean one, then births from (*w,n*) to (*w* + *z_i_*, *n* + 1) to are attempted at times 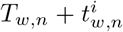 and where *T_w_,_n_* is the time (*l, m, n*) is first occupied. The births succeed if the sites are not yet occupied. Figure 7 show cross sections of the the growing tumor at *n* = 400 and *n* = 2000. Note that in contrast to the voter model simulation in Figure 6, the regions occupied by different types are solid, as we would expect in a growing tumor.

**Figure 7:**
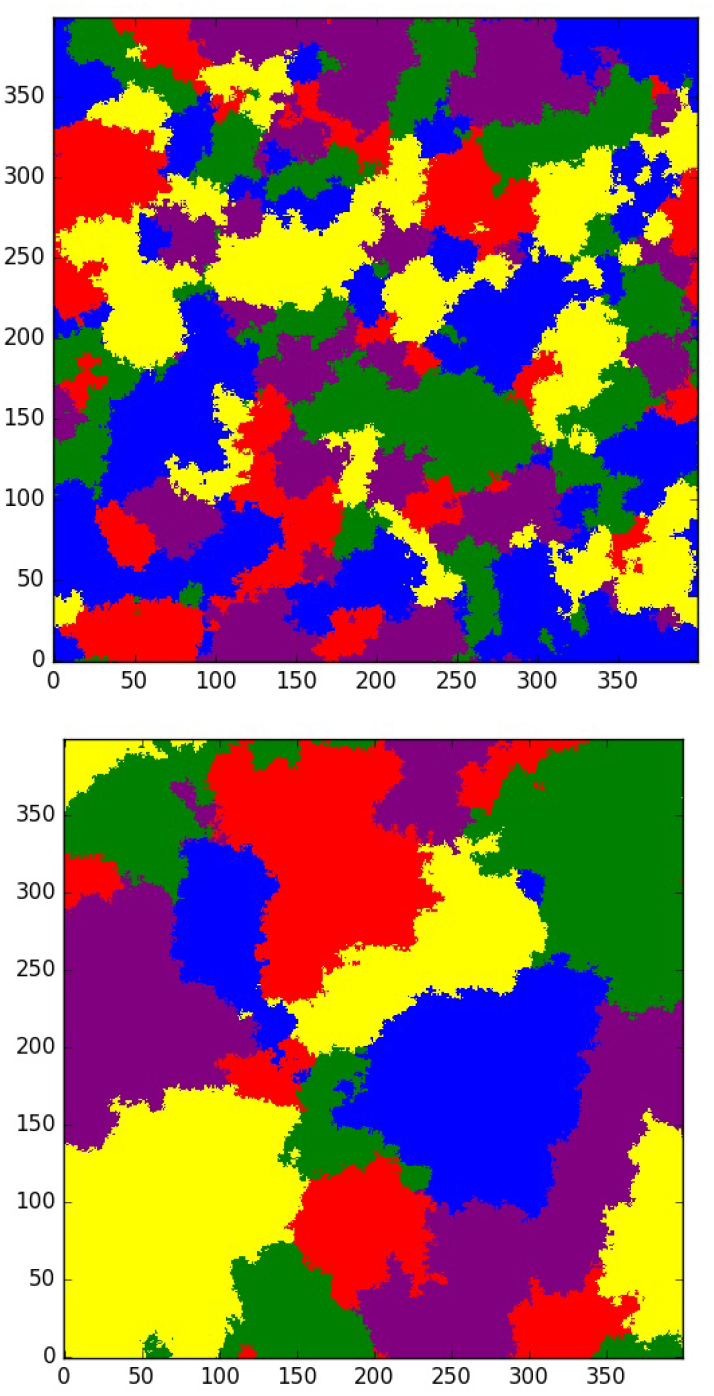
Two dimensional contact process at times 400 and 2000.

Fusco et al [16], see their Table 1, predicted that clone sizes in three dimensions has 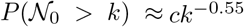. Figure gives a log-log plot of the number of types versus time in a 400 × 400 simulation. The slope is ¾ 1.32. Computing as before

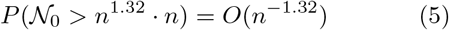

Changing variables *n* = *k*^1/2.32^, gives 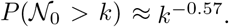.

## 6 Conclusions

We have investigated two models for the growth of a solid tumor with the aim of reproducing genotype patterns found in a recent study of colon cancer [34]. In both models growth only occurs at the boundary. In two dimensions both the voter and contact versions of the model produce solid regions of cells that have the same genotype. However, the two models make different predictions about the sizes of these regions. In three dimensions, the voter model has unrealistic small scale heterogeneity, while the contact process version produces solid regions with smooth boundaries.

Based on the results presented here it seems that the contact process model, which is equivalent to first passage percolation reproduces genotype patterns found in solid tumors. Genealogies are not easily to study mathematically. Boundary fluctuations in two dimensions are conjectured to have a complicated limit distribution. In *d* = 3 we have no idea how to compute the probability that a type survives to time *n*. However, the process is easily simulated and the observed patterns can be compared to sequencing data from cancer tumors.

**Figure 8:**
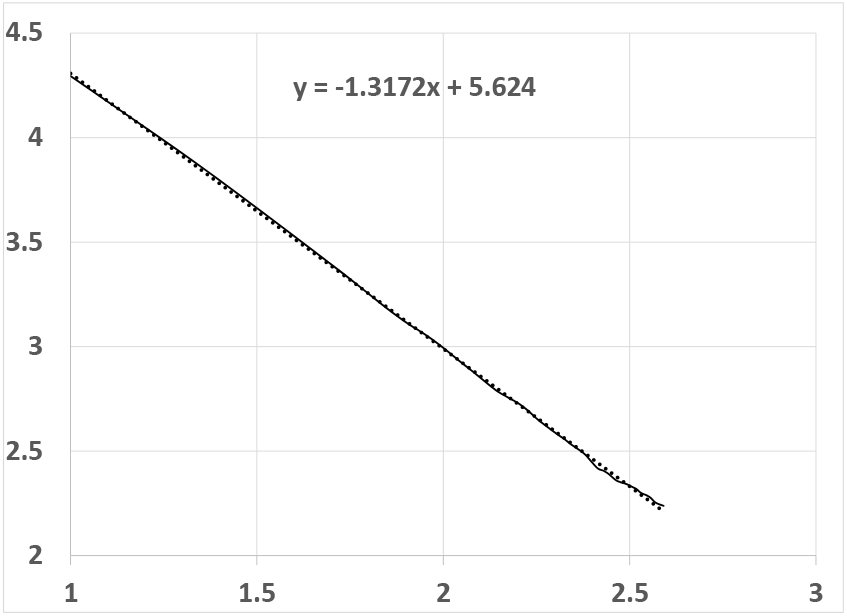
Log-log pot of the number of regions versus time in the 2 + 1 dimensional contact process.

## A Proof of Theorem 1

Let *B_t_*, *t* ≥ 0 be Brownian motion and let *τ*_0_ = inf {t : *B_t_* = 0 }. Let 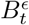 be a Brownian motion started at ∈ > 0 and let

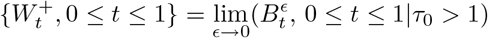

be the Brownian meander. In [11] it was shown that if *S_n_* is a simple random walk with *S*_0_ = 1 then

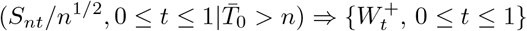

where the weak convergence occurs in C[0,1]. Using the continuous mapping theorem it follows that

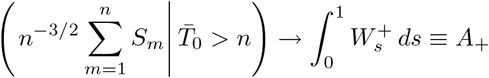

The corresponding result for our lazy random walk is

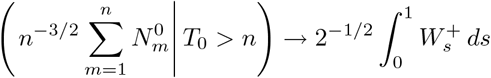

Combining this with (1) gives a limit result for the size of families conditioned that they do not die out by time *n*

The Brownian excursion 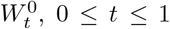, 0 ≥ *t* ≥ 1 can be defined as either the Brownian Bridge conditioned to stay positive or the Brownian meander conditioned to end at 0. It is known that if Sn is a simple random walk with *S*_0_ = 1 then as *n* → ∞ through the even integers

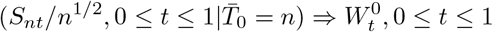

where the weak convergence occurs in *C*[0,1]. Using the continuous mapping theorem it follows that

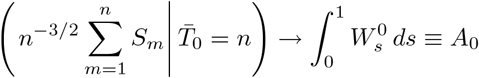

The corresponding result for the lazy random walk is

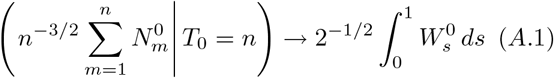

The area of the Brownian excursion, *A*_0_, arises in the enumeration of connected graphs [41, 35] and many other problems in combinatorial theory [22]. Takacs [37] found an explicit formula as an infinite series of confluent hypergeometric functions. Janson and Louchard [23] derived asymptotics for the density

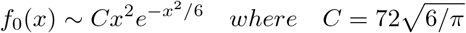

Janson [22] computed moments of the Brownian excursion 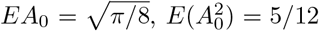 and the Brownian meander.

Let 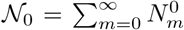. Combining (1) with (A.1) we see that

### Theorem 2.

*As* 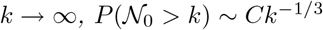

*Proof.* Let 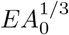. Replacing the sum by an integral

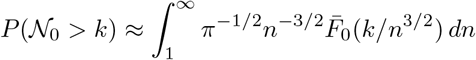

Changing variables *y* = *kn*^−3/2^, i.e., *n* = *k*^2/3^ *y*^2/3^ and *dn* = (−2/3)*k*^2/3^*y*^−5/3^ *dy* we have

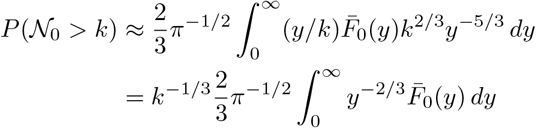

The integral is 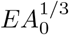.

## References

[1] Auffinger, A., Damron, M., and Hanson, J. (2016) 50 years of first passage percolation. arXiv: 1511.03262

[2] Bozic, I. et al. (2010) Accumulation of driver and passenger mutations during tumor progression. Proc. Natl. Acad. Sci. 107(43), 18545–18550

[3] Bozic, I. et al (2013) Evolutionary dynamics of cancer in response to targeted combination therapy. eLife 2, e00747.

[4] Bozic, I., Gerold, J.M., and Nowak, M.A. (2016) Quantifying clonal and subclonal passenger mutations in cancer evolution. PLOS Computational Biology 12, e1004731.

[5] Bramson, M., and Griffeath, D. (1980) Asymptotics for Interacting Particle Systems on Z*^d^*. Z. Wahrscheinlichkeitstheorie verw. gebiete. 53, 183–196

[6] Bramson, M., and Griffeath, D. (1980) On the Williams-Bjerknes tumour growth model. II. Math. Proc. Cambridge Philos. Soc. 88, 339–357

[7] Bramson, M., and Griffeath, D. (1981) On the Williams-Bjerknes tumour growth model. I. Ann. Probab. 9, 173–185

[8] Cox, J.T., and Griffeath, D. (1986) Diffusive clustering in the two dimensional voter model. Annals of Probability. 14, 347–370

[9] Derrida, B. and Dickman, R. (1991). On the interface between two growing Eden clusters. J. Physics A., 24, 1385–1422

[10] Ding, Li, et al. (2012) Clonal evolution in relapsed acute myeloid leukemia revealed by whole-genome sequencing. Nature. 481, 506–510

[11] Durrett, R. (1978) Conditional limit theorems for some null recurrent Markov processes. Ann. Probab. 6, 798–828

[12] Durrett, R. (2015) Branching Process Models of Cancer. Springer-Verlag, New York

[13] Durrett, R. (2017) Genealogies in growing solid tumors. bioRxiv 244160

[14] Durrett, R., Foo, J., Leder, K., Mayberry, J., Michor, F. (2011) Intratumor heterogeneity in evolutionary models of tumor progresssion. Genetics. 188, 461–477

[15] Eden, Murray (1961) A two-dimensional growth process. Pages 223–239 in Vol IV of Proc. 4th Berkeley Sympos. Math. Statist. and Prob. U. California Press, Berkeley, Calif.

[16] Fusco, D., Gralka, M., Kayser, J., Anderson, A., and Hallatschek, O. (2016) Excess of mutational jackpot events in expanding populations revealed by spatial Luria-Delbrück experiments Nature Communications. 7, paper 12760

[17] Gerlinger, M, et al (2015) Intratumor heterogeneiry and branched evolution revealed by multiregion seqeuncing. The New England Journal of Medicine. 366, 883–892

[18] Hallatschek, O., Hersen, P., Ramanathan, S., and Nelson, D.R. (2007) Genetic drift at expanding frontiers promotes gene segregation. Proc. Natl. Acad. Sci. 104, 19926–19930

[19] Hallatshek, O., and Nelson, D.R. (2008) Gene surfing in expanding populations. Theor. Pop. Biol. 73, 158–170

[20] Hallatschek, O., and Nelson, D.R. (2009) Life at the front of an expanding population. Evolution. 64, 193–206

[21] Iwasa, Y., Nowak, M.A., and Michor, F. (2006) Evolution of resistance during clonal expansion. Genetics. 172, 2557–2566

[22] Janson, S. (2007) Brownian excursion area, Wright’s constants in graph enumeration, and other Brownian areas. Probability Surveys. 4: 80145.

[23] Janson, S., and Louchard, G. (2007) Tail estimates for the Brownian excursion area and other Brownian areas. Electronic Journal of Probability. 12: 16001632

[24] Kardar, M., Parisi, G., and Zhang, Y.C. (1986) Dynamic scaling of growing interfaces. Phys. Rev. Letters. 56, 889–892

[25] Kaveh K, Takahashi Y, Farrar MA, Storme G, Guido M, Piepenburg J, Penning J, Foo J, Leder KZ, Hui SK. (2017) Combination therapeutics of Nilotinib and radiation in acute lymphoblastic leukemia as an effective method against drug-resistance. PLoS Computational Biology. 13(7):e1005482.

[26] Korolev, K.S. et al. (2012) Selective sweeps in growing microbial colonies. Physical Biology. 9, paper 026008

[27] Leder, K., Foo, J., Skaggs, B., Gorre, M., Sawyers, C.L., and Michor, F. (2011) Fitness conferred by BCR-ABL kinase domain mutations determines the risk of pre-existing resistance in chronic myeloid leukemia. PLoS One. 6, paper e27682

[28] Mumenthaler S, Foo J, Leder K, et al. (2011) Evolutionary modeling of combination strategies to overcome resistance to tyrosine kinase inhibitors in non-small cell lung cancer. Mol. Pharm. (2011) 8(6).

[29] Navin, N., et al (2011) Tumour evolution inferred by single-cell sequencing. Nature. 472, 90–94

[30] Richardson, D. (1973) Random growth in a tesselation. Proc. Camb. Phil. Soc. 74, 515–528

[31] Ryser, M.D., Min, B.H., Siegmund, K., and Shibata, D. (2017) Spatial mutaiton patterns as markers of early tumor cell mobility. Proc. Natl. Acad. Sci. 115, 5774–5779

[32] Saito, Y., and Müller-Krumbhaar, H. (1995) Critical phenomena in morphology transitions of growth models with competition. Phys. Rev. Letters. 74, 4325–4328

[33] Sottoriva, A., et al. (2013) Intratumor heterogeneity in human glioblastoma reflects cancer evolutionary dynamics. Proc. Natl. Acad. Sci. 110, 4009–4014

[34] Sottoriva, A., et al. (2015) A Big Bang model of human colorectal cancer growth. Nature Genetics. 47, 209–216

[35] Spencer, J. (1997) Enumerating graphs and Brownian motion. Comm. Pure Appl. Math. 50, 291294.

[36] Sun, R., Hu, Z., and Curtis, C. (2018) Big bang tumor growth and clonal evolution. Cold Spring Harbor Perspectives in Medicine. 8: a028381

[37] Tackacs, L. (1991) A Bernoulli Excursion and its various applications. Adv. Appl. Probab. 23, 557–585

[38] Tomasetti, C., and Levy, D. (2010) Roles of symmetric and asymmetric division of stem cells in the evolution of resistance. Proc. Natl. Acad. Sci. 107, 16766–16771

[39] Tomasetti, C., Vogelstein, B., and Parmigiani, G. (2013) Half or more somatic mutations in cancers of self-renewing tissues originate prior to cancer initiation. Proc. Natl. Acad. Sci. 110, 1999–204

[40] Williams, T., and Bjerknes, R. (1972) Stochastic model for abnormal clone spread through epithelial basal layer. Nature. 235, 19–21

[41] Wright, E.M. (1977) The number of connected sparsely edged graphs. J. Graph Theory. 1, 317–330

[42] Yachida, S., et al (2010) Distant metastases occur late during the genetic evolution of pancreatic cancer. Nature. 467, 1114–1117

